# Semi-quantitative characterisation of mixed pollen samples using MinION sequencing and Reverse Metagenomics (RevMet)

**DOI:** 10.1101/551960

**Authors:** Ned Peel, Lynn V. Dicks, Darren Heavens, Lawrence Percival-Alwyn, Chris Cooper, Matthew D. Clark, Richard G. Davies, Richard M. Leggett, Douglas W. Yu

## Abstract

1. The ability to identify and quantify the constituent plant species that make up a mixed-species sample of pollen has important applications in ecology, conservation, and agriculture. Recently, metabarcoding protocols have been developed for pollen that can identify constituent plant species, but there are strong reasons to doubt that metabarcoding can accurately quantify their relative abundances. A PCR-free, shotgun metagenomics approach has greater potential for accurately quantifying species relative abundances, but applying metagenomics to eukaryotes is challenging due to low numbers of reference genomes.

2. We have developed a pipeline, RevMet (Reverse Metagenomics), that allows reliable and semi-quantitative characterization of the species composition of mixed-species eukaryote samples, such as bee-collected pollen, without requiring reference genomes. Instead, reference species are represented only by ‘genome skims’: low-cost, low-coverage, short-read sequence datasets. The skims are mapped to individual long reads sequenced from mixed-species samples using the MinION, a portable nanopore sequencing device, and each long read is uniquely assigned to a plant species.

3. We genome-skimmed 49 wild UK plant species, validated our pipeline with mock DNA mixtures of known composition, and then applied RevMet to pollen loads collected from wild bees. We demonstrate that RevMet can identify plant species present in mixed-species samples at proportions of DNA ≥1%, with few false positives and false negatives, and reliably differentiate species represented by high versus low amounts of DNA in a sample.

4. The RevMet pipeline could readily be adapted to generate semi-quantitative datasets for a wide range of mixed eukaryote samples, which could include characterising diets, quantifying allergenic pollen from air samples, quantifying soil fauna, and identifying the compositions of algal and diatom communities. Our per-sample costs were £90 per genome skim and £60 per pollen sample, and new versions of sequencers available now will further reduce these costs.

## Introduction

Pollination is a key ecosystem service; almost 90% of all flowering plant species, including 75% of food crops (mainly fruits, nuts, and vegetables), rely on animal pollination (Ollerton, Winfree, & Tarrant, 2011; Klein *et al.,* 2007). The benefits of pollinators, and pollinator-dependent plants, also include the production of medicines, biofuels, fibres, and construction materials (Potts *et al.,* 2016). There is growing concern over the decline of wild and domesticated pollinators and the resulting decrease in pollination services and crop production (Potts *et al.,* 2010; Burkle, Marlin, & Knight, 2013). These declines are thought to be caused by multiple threats acting together, including habitat loss, climate change, and the spread of diseases (Vanbergen *et al.,* 2013).

To mitigate drivers of pollinator decline, the Intergovernmental Science – Policy Platform for Biodiversity and Ecosystem Services (IPBES) has suggested three complementary strategies: (1) ecological intensification, which involves boosting agricultural production by increasing the provision of supporting ecological processes such as biotic pest regulation, nutrient cycling, and pollination (Bommarco, Kleijn, & Potts, 2013; Tittonell, 2014); (2) strengthening existing diversified farming systems, including gardens and agroforestry, for the generation of ecosystem functions; and (3) investment in ecological infrastructure, to protect, restore, and connect natural and semi-natural habitats across agricultural landscapes, so that pollinator species can more easily disperse and find nesting and floral resources (IPBES 2016).

However, knowledge gaps limit the effectiveness of these strategies (Wood, Holland, & Goulson, 2015; Dicks *et al.,* 2013). For instance, it is still not clear which plant species are the most valuable food resources and how plant species vary in value across pollinator species, over time, and in different environmental conditions. It is also not well understood whether the addition of floral resources might draw pollinators away from pollinator-dependent crop plants (Morandin & Kremen, 2013), or whether floral enhancement will alter levels of plant-target specialism, at the levels of insect species and of individual insects, resulting in changes in pollination efficiency (Lucas *et al.*, 2018; Morales & Traveset, 2008).

Therefore, a crucial technical challenge for understanding plant-pollinator interactions is a method to identify *and* quantify the species of pollen that are consumed by pollinators. Identifying and quantifying pollen has traditionally been carried out by using light microscopy to distinguish plant species by grain morphology, a labour-intensive technique that requires expert knowledge and lacks discriminatory power at lower taxonomic levels (Long & Krupke, 2016; Khansari *et al*., 2012). In contrast, high-throughput DNA sequencing now allows pollen identification without expert knowledge of pollen morphology and taxonomy.

The currently dominant sequence-based method is metabarcoding, which involves amplifying taxonomically informative marker genes from mixed samples via polymerase chain reaction (PCR) (Ji *et al.*, 2013). The resulting amplification products, known as amplicons, are sequenced, and the reads are assigned to taxonomies by matching against barcode databases, such as the Barcode of Life Data System (Ratnasingham & Hebert, 2007). Notably for plants, there is no single barcode gene that matches the resolving power and universality of 16S rRNA for prokaryotes and Cytochrome Oxidase (CO1) for animals (Hollingsworth, Li, Van Der Bank, & Twyford, 2016). Instead, plant-related barcoding studies rely on a combination of marker genes, which include plastid regions *rbcL* and *matK* and the internal transcribed spacer (ITS) regions of nuclear ribosomal DNA (Li *et al.,* 2015; Hollingsworth *et al.,* 2016). Metabarcoding of mixed-species pollen samples can reveal the presence and absence of constituent plant species (or genera), but there are strong reasons to doubt that metabarcoding can accurately quantify their relative abundances, due to PCR amplification biases and varying copy numbers of barcode loci (Keller *et al.,* 2015; Richardson *et al.,* 2015; Sickel *et al.,* 2015; Bell *et al.,* 2017, 2018; Lamb et al. 2018).

In contrast to the targeted sequencing approach of metabarcoding, ‘shotgun metagenomics’ involves randomly sequencing short stretches of genomic DNA from mixed samples. In standard metagenomics, these short reads (‘queries’) are mapped to either assembled genomes or to collections of barcode genes (‘references’), which creates a requirement for large numbers of reference genomes (Sharpton, 2014) or barcodes, with the latter being very inefficient. Species identification is obtained by first calculating a similarity metric between each short read and each reference sequence (e.g. % identity) and then using an algorithm to assign each short read to the most likely reference sequence (Quince, Walker, Simpson, Loman, & Segata, 2017). The potential key advantages of shotgun metagenomics are that it can avoid the PCR-induced biases seen with metabarcoding, especially if PCR-free library preparation protocols are used (see Nayfach & Pollard, 2016; Jones *et al.,* 2015) and that by sampling across the whole genome, variation in the copy numbers of a few loci is rendered less important. However, the requirement for reference genomes means that most shotgun metagenomics studies focus on prokaryotic organisms, since large numbers of prokaryote reference genomes are available. In contrast, eukaryotes are not well represented in sequence databases and as a result have mostly been neglected in metagenomic studies (Escobar-Zepeda, De León, & Sanchez-Flores, 2015). The low numbers of reference genomes for eukaryotic species is because they are more expensive to sequence and assemble (Gilbert & Dupont, 2011).

Here we demonstrate a metagenomic pipeline for eukaryotes that avoids the need to assemble reference genomes. Instead, each reference species is represented by a ‘genome skim,’ which is a low-cost, low-coverage, shotgun dataset, i.e. simply a set of short reads. These sets of short reads are used to identify individual *long reads* from pollen that have been generated by sequencing mixed-species pollen loads with the Oxford Nanopore Technologies’ (ONT) MinION, a nanopore sequencing device (for a review of MinION applications and performance, see Leggett & Clark, 2017). Here, we generate reference genome skims for 49 wild UK plant species, and we use them to identify and quantify plant species in two kinds of query samples: mock, mixed-plant-species DNA mixtures of known composition and mixed-species pollen samples collected from wild bees. Since the long reads in each query sample are individually identified, we show that the frequency of long reads assigned to a plant species is a reasonably accurate estimate of that species’ biomass frequency in a mixed-species sample. We call this pipeline Reverse Metagenomics, or RevMet, because we map reference sequences to query sequences, which is the reverse of the normal metagenomic protocol.

## Methods

### Sampling of bees and plant tissue

Sample collection took place in the Pensthorpe Natural Park area (52°49’23’’N, 0°53’14’’E) of Norfolk, UK, during June and July 2016. Leaf samples were collected from all plant species with open flowers, including grasses and trees, within a 100 m radius of the collection site (n = 49 species). Leaf tissue was preserved on dry ice in the field followed by storage at −80 °C. Foraging wild bees (n = 48: 9 *Apis mellifera*, 27 *Bombus terrestris/lucorum complex*, 12 *Bombus lapidarius*) were collected with hand nets or into falcon tubes directly from flowers and euthanized in falcon tubes containing ethanol-soaked tissue paper. Pollen loads were scraped from bee corbiculae using a mounted needle and stored in absolute ethanol. The plant species on which each bee was foraging when collected was recorded.

### Leaf tissue DNA extraction, library preparation, and Illumina sequencing

Leaf tissue from each of the 49 plant species was disrupted by bead-beating using a 4-mm stainless steel bead with a Qiagen TissueLyser II running at 22.5 Hz for 4 min, rotating the adapter sets after 2 min. DNA was extracted using the DNeasy Plant Kit (Qiagen, Hilden, Germany) following manufacturer’s instructions. DNA concentrations were measured on a Qubit 2.0 fluorometer (ThermoFisher, Waltham, USA) using the dsDNA HS assay kit, and fragment size distribution was checked with a Genomic DNA Analysis ScreenTape on the TapeStation 2200 (Agilent, Santa Clara, USA).

The Earlham Institute (Norwich, UK) applied a modified version of Illumina’s Nextera protocol, known as Low Input Transposase Enabled (LITE) protocol (Beier *et al.,* 2017), to generate a separate sequencing library for each leaf sample, targeting an average insert size of 500 bp. The LITE libraries were then pooled based on estimated genome sizes (Supplementary Table S1), obtained from the Royal Botanic Gardens Kew Plant DNA C-values database (Bennett and Leitch, 2012), in order to achieve 0.5x coverage of each species genome. The pooled libraries were sequenced on one lane of Illumina HiSeq 2500 in Rapid Run mode (250 bp PE).

### Construction and sequencing of mock pollen samples

DNA from twelve of the 49 plant species were used to construct six mock communities. Each mock was made using 200 ng DNA in total, with species added at different proportions: 0.08% to 45.25% (Table 1). For each mock, technical-replicate pairs were prepared using ONT’s (Oxford, UK) Rapid Barcoding Sequencing Kit (SQK-RBK001), following the RBK_9031_v2_revl_09Mar2017 version of the manufacturer’s protocol. The 12 libraries (six mocks, duplicated) were sequenced on a single MinION R9.5 flow cell (FLO-MIN107).

**Table 1.** DNA mock mix compositions.

|  | *Knautia*<br>*arvensis* | *Galium*<br>*verum* | *Crepis*<br>*capillaris* | *Papaver*<br>*somniferum* | *Anagallis*<br>*arvensis* | *Sambucus*<br>*nigra* | *Bryonia*<br>*dioica* | *Ranunculus*<br>*repens* | *Lotus*<br>*corniculatus* | *Digitalis*<br>*purpurea* | *Leucanthemum*<br>*vulgare* | *Stachys*<br>*sylvatica* |
| --- | --- | --- | --- | --- | --- | --- | --- | --- | --- | --- | --- | --- |
| MM1.Ratios | 100 | 100 | 100 | 10 | 10 | 10 | 1 | 1 | 1 | 0 | 0 | 0 |
| MM2.Ratios | 0 | 0 | 0 | 1 | 1 | 1 | 10 | 10 | 10 | 100 | 100 | 100 |
| MM3.Ratios | 0 | 0 | 0 | 0 | 0 | 0 | 10 | 100 | 0 | 0 | 0 | 0 |
| MM4.Ratios | 0 | 0 | 0 | 1 | 0 | 1000 | 0 | 0 | 100 | 0 | 0 | 100 |
| MM5.Ratios | 100 | 100 | 100 | 0 | 1 | 1 | 1 | 100 | 0 | 0 | 0 | 1 |
| MM6.Ratios | 1 | 1 | 1 | 100 | 100 | 100 | 100 | 0 | 0 | 0 | 0 | 1 |
| MM1.DNA (ng) | 60.1 | 60.1 | 60.1 | 6.0 | 6.0 | 6.0 | 0.6 | 0.6 | 0.6 | 0.0 | 0.0 | 0.0 |
| MM2.DNA (ng) | 0.0 | 0.0 | 0.0 | 0.6 | 0.6 | 0.6 | 6.0 | 6.0 | 6.0 | 60.1 | 60.1 | 60.1 |
| MM3.DNA (ng) | 0.0 | 0.0 | 0.0 | 0.0 | 0.0 | 0.0 | 9.1 | 90.5 | 0.0 | 9.1 | 90.5 | 0.9 |
| MM4.DNA (ng) | 0.0 | 0.0 | 0.0 | 0.2 | 0.0 | 166.5 | 0.0 | 0.0 | 16.7 | 0.0 | 0.0 | 16.7 |
| MM5.DNA (ng) | 49.5 | 49.5 | 49.5 | 0.0 | 0.5 | 0.5 | 0.5 | 49.5 | 0.0 | 0.0 | 0.0 | 0.5 |
| MM6.DNA (ng) | 0.5 | 0.5 | 0.5 | 49.5 | 49.5 | 49.5 | 49.5 | 0.0 | 0.0 | 0.0 | 0.0 | 0.5 |
| MM1.Percentages | 30.0% | 30.0% | 30.0% | 3.0% | 3.0% | 3.0% | 0.3% | 0.3% | 0.3% | 0.0% | 0.0% | 0.0% |
| MM2.Percentages | 0.0% | 0.0% | 0.0% | 0.3% | 0.3% | 0.3% | 3.0% | 3.0% | 3.0% | 30.0% | 30.0% | 30.0% |
| MM3.Percentages | 0.0% | 0.0% | 0.0% | 0.0% | 0.0% | 0.0% | 4.5% | 45.3% | 0.0% | 4.5% | 45.3% | 0.5% |
| MM4.Percentages | 0.0% | 0.0% | 0.0% | 0.1% | 0.0% | 83.3% | 0.0% | 0.0% | 8.3% | 0.0% | 0.0% | 8.3% |
| MM5.Percentages | 24.8% | 24.8% | 24.8% | 0.0% | 0.3% | 0.3% | 0.3% | 24.8% | 0.0% | 0.0% | 0.0% | 0.3% |
| MM6.Percentages | 0.3% | 0.3% | 0.3% | 24.8% | 24.8% | 24.8% | 24.8% | 0.0% | 0.0% | 0.0% | 0.0% | 0.3% |

### Bee-collected pollen DNA extraction, library preparation, and MinION sequencing

After removing storage ethanol from the 48 bee-collected pollen loads, the pollen was disrupted with ca. five 1-mm stainless steel beads for 2 min at 22.5 Hz using a Qiagen TissueLyser II, rotating the adapter sets after 1 min. The pollen samples were resuspended in 600 μl CTAB extraction buffer (2% CTAB, 1.4 M NaCl, 20 mM EDTA, pH 8.0, 100 mM Tris-HCl pH 8.0), 0.5 μl of β-Mercaptoethanol, 4 μl of proteinase K, and vortexed for 5 s. Following a 1 hr incubation at 55 °C, the tubes were centrifuged for 6 min at 18,000 × g. The ≈500 μl of supernatant was extracted to a clean 1.5 ml tube before an equal volume of chilled (2-8 °C) Phenol:Chloroform:Isoamyl Alcohol (25:24:1, v/v) was added to the lysate. The samples were vortexed for 10 s (5 × 2 s bursts), centrifuged for 5 min at 14,000 × g, and the upper aqueous phase (≈ 420 μl) was extracted by pipette and transferred into a clean 1.5 ml tube.

An equal volume of Agencourt AMPure XP beads was added to each sample, vortexed for 20 s (10 × 2 s bursts), and then incubated for 10 min at room temperature. By placing the samples onto a magnetic tube rack for 5 min, the beads were separated from the solution, and the cleared supernatant was removed by aspiration. The beads were washed twice using the following protocol: 1 ml of 80% ethanol was added, incubated at room temperature for 30 s, and then removed, followed by air drying for ≈3 min. The magnetic beads were resuspended in 55 μl of EB (Elution Buffer: 10 mM Tris-HCl) and incubated at 37 °C for 10 min. The tubes were placed back onto the magnetic rack to bind the beads, and the eluted DNA (≈ 50 μl) was transferred into fresh tubes. A 1 μl aliquot of 1-in-10 diluted Qiagen RNase A was added to each DNA sample before being incubated for 30 min at 37 °C. The concentration of the eluted DNA was assessed using the dsDNA HS assay on a Qubit 2.0 fluorometer. To check the DNA for degradation, fragment size distributions were checked with a TapeStation 2200 using the Genomic DNA Analysis ScreenTape.

Finally, the extracted DNA was prepared and sequenced using the same protocol as used for the DNA mocks above, except that only one library was prepared for each sample. Twelve samples can be multiplexed using the Rapid Barcoding Sequencing Kit; we thus required four flow cells. Due to continuous software upgrades by ONT, the specific software versions of *MinKNOW* varied across runs and is recorded in the final sequence files (fast5 format), which are available from the EBI’s European Nucleotide Archive (see Data accessibility).

### Illumina and MinION read pre-processing

Duplicate reads were removed from the 49 plant-reference Illumina datasets using *NextClip* (Leggett, Clavijo, Clissold, Clark, & Caccamo, 2014), and then *cutadapt 1.10* (Martin, 2011) was used to trim Illumina adaptors and filter out reads shorter than 100 bp. The resulting unmerged FASTQ files constitute our 49 *reference skims*.

The MinION datasets from the 12 mocks and the 48 pollen loads were basecalled and demultiplexed with *albacore 2.1.10* (ONT). The resulting FASTQ files were converted to FASTA format. We removed long reads deriving from plant organelles because they are highly conserved across plant species and in pilot tests we observed that mapping to organellar long reads resulted in a higher rate of incorrect assignments than mapping to nuclear long reads (data not shown). NCBI Entrez (https://www.ncbi.nlm.nih.gov/sites/batchentrez) was used to download 2,583 Land Plant organelle genomes, including 1,852 chloroplast, 226 mitochondrial, and 505 plastid genomes. Organelle reads were identified by aligning each of the MinION datasets to the organellar genomes using *minimap2 2.7* (Li, 2018) and removed from the FASTA files. The resulting 60 (= 12 + 48) organelle-filtered FASTA files constitute our mock and pollen *query* datasets, and in the next step, we used the 49 plant reference skims to assign a taxonomy to each long read in the mock and pollen query datasets (Fig. 1c).

**Figure 1.**
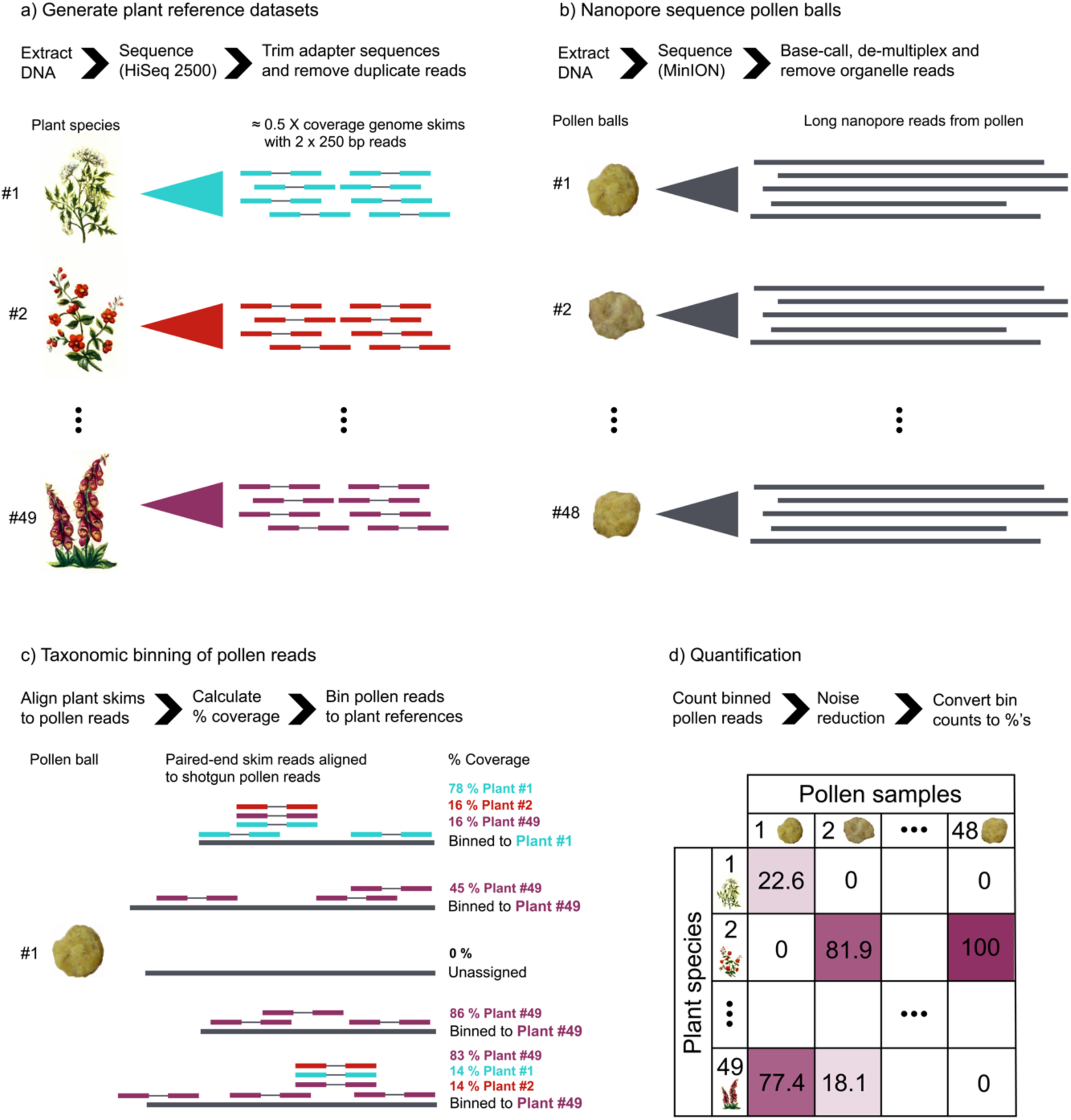
RevMet pipeline overview. a) Low coverage, short-read, reference datasets were generated for 49 wild plant species. b) Bee-collected pollen loads were sequenced on a MinION, generating long read datasets. c) The 49 short-read reference datasets were separately mapped to the long-read pollen datasets, and each pollen read was assigned to the plant species that mapped with the highest percent coverage or was left unassigned if the highest coverage was <15%. d) Binned pollen reads were counted, noise was reduced by implementing a 1% minimum-abundance filter, and then the remaining bin counts were converted to percentages.

### Taxonomic assignment of mock-sample and bee-collected pollen MinION reads

We used *bwa mem 0.7.17* (Li, 2013) to map the Illumina reads from each of the 49 reference skims against every individual long read in each of the mock and bee-collected pollen datasets. *SAMtools 1.7* (Li *et al.,* 2009) was used to remove unmapped reads and secondary and supplementary alignments. After SAMtools indexing, the depth of mapping coverage at each long-read position was calculated using the SAMtools *depth* function, followed by the calculation of ‘percent coverage’ for each long read (using a custom python script), defined as the fraction of nucleotide positions that were mapped to by one or more reference-skim Illumina reads.

We assigned each long read to the plant species that mapped with the highest percentage coverage, unless the highest percent coverage was <15%, in which case the long read’s identity was judged ambiguous and left unassigned. Additionally, for clarity of presentation, we implemented a 1% minimum-abundance filter, removing plant species represented by fewer than 1% of the total assigned long reads in each sample.

### Reference-skim subsampling

To estimate a minimum recommended depth of coverage needed per reference skim, we subsampled one of the genome skims, *Knautia arvensis*, which is a major constituent species in mock mixes MM1 and MM2. We randomly subsampled this skim from its maximum of 0.65x down to 0.05x, in steps of 0.05x using a custom script. For each subsample, the whole pipeline was re-run along with the full reference skims of the other 48 plant species. The number of mock reads assigned to *Knautia arvensis*, and the number of unassigned reads, at each level of coverage was recorded. This subsampling was repeated three times (Supplementary Fig. S1).

### Network construction

We constructed a pollinator-plant network diagram for the 48 wild-bee pollen samples, using the *bipartite 2.11* package (Dormann, Frund, Bluthgen, & Gruber, 2009) for the *R* statistical language (R Core Team, 2018). For presentational presentational clarity, we only show plant species represented by more than 10% of the assigned reads in each sample.

## Results

### A reference set of plant genome skims

Low genome-coverage, short-read, shotgun-sequencing datasets (‘reference skims’) were successfully generated for all 49 plant species (Fig. 1a). After pre-processing, the mean estimated coverage was 0.6x (0.1 to 1x, details in Supplementary Table S1).

### Mock DNA mixes

The six mock communities, each with two technical replicates, were sequenced on a MinION. These produced reads with mean length 1914 bp (longest 41,058 bp). After demultiplexing, 88.8% of the reads could be assigned to one of the 12 mock mixes, with the remaining reads left unclassified. Sequences originating from organellar genomes made up between 5.1% (MM4.2) to 10.2% (MM3.2) of the reads in the mocks and were removed. The remaining number of reads per mock ranged from 733 (MM2.1) to 2174 (MM4.1), mean 1347.

### Taxonomic assignment of mock-sample MinION reads

The 49 reference skims were separately mapped to each long read in each of the 12 mock mixes, and each long read was assigned to the plant species that mapped with the highest percent coverage, or left unassigned if the highest coverage was <15%. In total, 65.5% of the mock reads were assigned to a plant species, with 94.7% of those reads being assigned to a species known to be present in that mock sample. Almost all (93.4%) of the 563 false-positive read assignments were made to one species, *Ranunculus acris*, and all these assignments occurred in the mock samples that contained the very closely related species *Ranunculus repens*. We return to this in the Discussion. The few other false-positive assignments all occurred at a rate of less than 1% of the assigned long reads in their mixes and for presentational clarity are not shown in Fig. 2. The full results are in Table S2.

**Figure 2.**
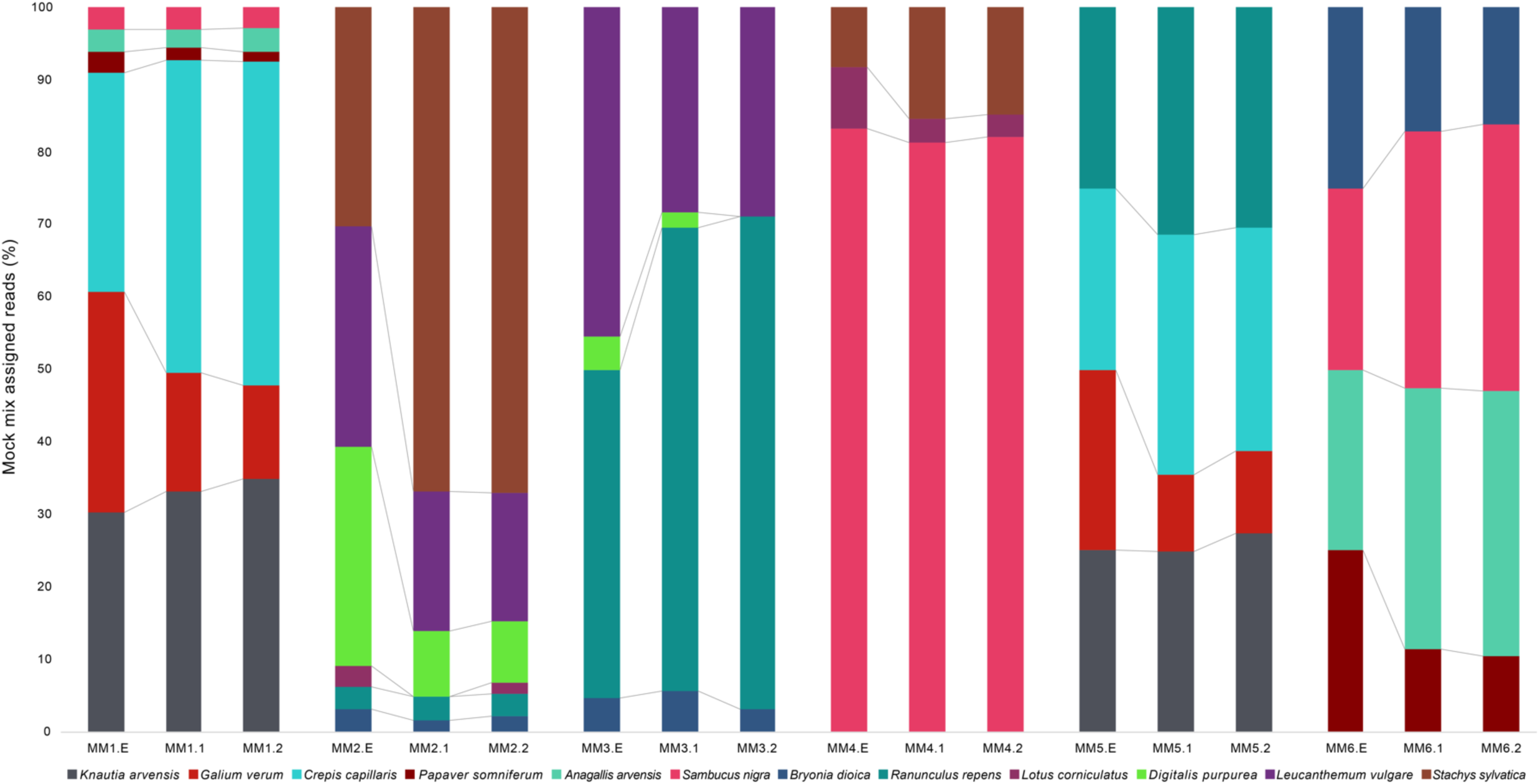
Expected vs observed mock mix compositions. Six mock plant DNA mixes, each with two technical replicates, were sequenced on a MinION and the RevMet method was applied. The first stacked bar of each triplet represents the expected proportions based on input DNA. The second and third bar of each triplet reflect the observed MinION read assignments resulting from this pipeline.

All of the plant species that had been added to the mock compositions at proportions ≥1% were detected by our method in at least one of the two replicates, and in general, the technical replicates showed a high level of repeatability (Fig. 2). In two of the mocks, there was one species each (*Lotus corniculatus* in MM2 and *Digitalis purpurea* in MM3) that were detected in only one of the two replicates. In these two replicates, *Lotus corniculatus* and *Digitalis purpurea* were expected to be present at only 3.0% and 4.6%, respectively. Both of these species were consistently underrepresented across our mock data sets.

### Reference-skim subsampling

As expected, the larger the reference-skim dataset size for *Knautia arvensis*, the more reads in the MM1 and MM2 mocks were assigned to this species and the fewer reads left unassigned. Importantly, the rate of increase was decelerating (Supplementary Fig. S1); over half of the MinION reads that were assigned to *Knautia arvensis* with a 0.65x genome skim could also be assigned with just a 0.1x skim, even though all the other reference skims in the mapping run were kept at their original sizes.

### Taxonomic assignment of bee-collected pollen MinION reads

The 48 bee-collected pollen loads harvested from the corbiculae of three species, *Apis mellifera, Bombus terrestris/lucorum* complex, and *Bombus lapidarius*, yielded DNA quantities ranging from 191 to 3750 ng, and all successfully produced libraries, demonstrating that pollen carried by individual bees can provide sufficient DNA for MinION sequencing.

As with the 12 mocks, each of the reference skims was aligned to each long read in each of the 48 pollen samples, the long reads were either assigned to the plant species achieving the highest percent coverage or left unassigned, and any plant species assigned fewer than 1% of the long reads in each bee-collected pollen sample was filtered out (Supplementary Table S3). In total, 49.7% of the long reads were assigned to one of the reference plant species. In 38 of the 48 bees (79.2%), pollen from the plant species on which each bee was captured was found to be present in that bee’s pollen load (Supplementary Table S3).

Each of the 48 pollen loads was found to contain one or two major plant species (defined as read frequency ≥10%) (Fig. 3a). All nine of the *Apis mellifera* pollen loads contained a single major species, whereas 16 of 27 *Bombus terrestris/locorum complex* and 6 of 12 *Bombus lapidarius* pollen loads were comprised of two major species (Fig. 3a). These differences in mean number of major species were statistically significant (*Apis mellifera* versus *Bombus terrestris/locorum complex* (Welch’s t-test, t = −6.15, df = 26, p-value < 0.0001) and versus *Bombus lapidarius* (t = −3.32, df = 11, p-value < 0.01)) (Fig. 3b). Another way of visualising the wild-bee results is as a plant-pollinator network graph (Fig. 3c). Overall, 6 of the 49 reference plant species were identified as major components in the 48 pollen loads, and the majority of bee-collected pollen samples were dominated by one plant species.

**Figure 3.**
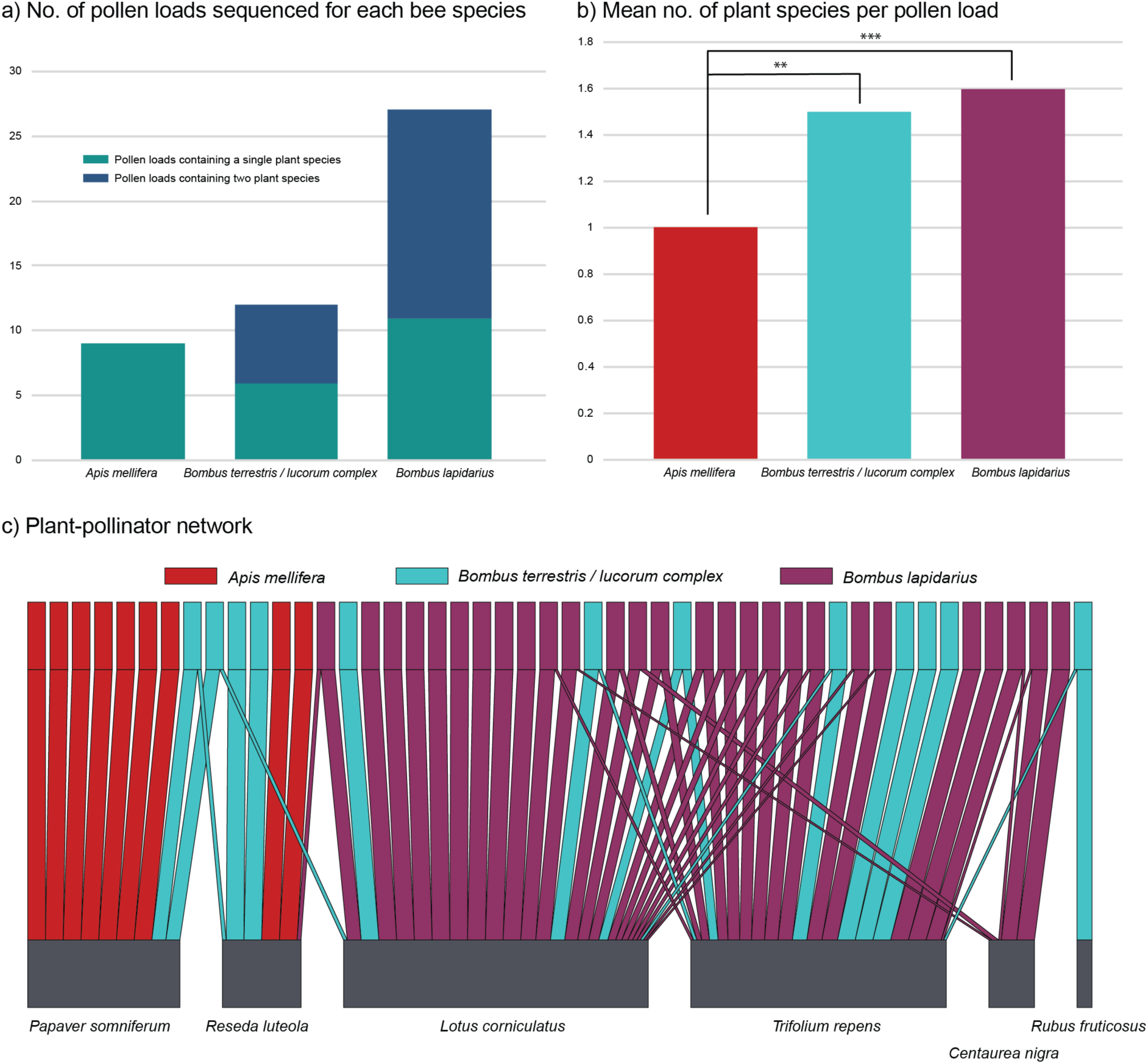
Bee-collected pollen compositions and plant-pollinator interactions. a) The number of individual pollen loads sequenced from three different species of bee. The proportion of pollen loads that contained a single major plant species are represented by green bars, while those with two major plant species are shown in blue. b) Mean number of plant species per pollen load for each of three different species of bee; ** p<0.01, *** p<0.001. c) Bipartite plant-pollinator network. The upper bars represent individual pollen loads from three different bee species, *Apis mellifera* (red), *Bombus terrestris/lucorum complex* (blue), and *Bombus lapidarius* (purple). The lower bars (grey) represent plant species. Link width indicates the MinION read proportion of each major plant species within each pollen load.

## Discussion

Using light microscopy to identify plant species from pollen requires expert knowledge and is costly when applied to many samples (Khansari *et al.,* 2012). There is a need for a quick and low-cost method that can be scaled to large numbers of pollen samples. Metabarcoding is the current leading candidate, but there are concerns over its discriminatory power at lower taxonomic levels, and there is good reason to believe that metabarcoding does not return reliable quantitative data (Keller *et al.,* 2015; Richardson *et al.,* 2015; Sickel *et al.,* 2015; Bell *et al.,* 2017, 2018; Lamb et al. 2018). A PCR-free shotgun-metagenomics approach has greater potential for providing reliable quantitative analysis with high power for resolving species. However, applying shotgun metagenomics to eukaryotes is challenging due to the lack of reference genomes (Gilbert & Dupont, 2011). We have developed a metagenomics method that avoids the need for reference genomes. Instead, each reference species is represented by just a low-cost genome skim, and we use a set of such skims to identify individual long reads from pollen samples, produced by the MinION sequencer.

We evaluated our RevMet pipeline with mock DNA mixtures of known composition and then applied the pipeline to pollen collected from wild bees. Our main findings are:

1. RevMet can identify plant species present in mixed-species samples at proportions of DNA ≥1%, with few false positives and false negatives, and can reliably differentiate species represented by high versus low amounts of DNA in a sample (Fig. 2, Supplementary Table S2).
2. Genome skims with sequence coverage as low as 0.05x can be used for detecting species presence and for estimating relative abundance in terms of DNA mass. Increasing skim coverage increases detection power, at a decelerating rate (Supplementary Fig. S1).
3. Individual pollen loads collected from wild *Apis* and *Bombus* bees yield enough DNA for MinION sequencing (Supplementary Table S3) and generate plausible plant-pollinator networks, as evidenced by the fact that (a) 56.3% of the plant species on which the bees were collected were also the dominant constituent of the corresponding pollen sample (and 79% of plant species on which the bees were collected were detected in the corresponding pollen sample) (Supplementary Table S3), and (b) pollen species richnesses and compositions were more similar within bee species than across bee species (Fig. 3).
4. Our per-plant-species cost of a reference skim was £90, and our per-pollen-sample cost was £61, including DNA extraction, library preparation, and sequencing. Sequencing costs will likely drop further, given the new Illumina NovaSeq and new MinION ‘Flongle’.

### Semi-quantitative species compositions

We were able to assign roughly 65% of the mock-mix MinION reads and just under 50% of the pollen-load MinION reads to our reference plant species. Importantly, the frequencies of MinION reads that were assigned to each reference plant species were reliably *‘semi-quantitative*,’ that is, able to differentiate low- and high-frequency plant species (Fig. 2). Within low- and high-abundance categories, accuracy was lower. For example, in mock sample MM1, *Knautia arvensis, Galium verum*, and *Crepis capillaris* were the three high-abundance species (each representing 30.3% of total input DNA mass each), and *Papaver somniferum, Anagallis arvensis*, and *Sambucus nigra* were the three low-abundance species (each representing 3.0% of total input DNA mass each). The RevMet pipeline estimated the three high-abundance frequencies at means of 34.0%, 14.7%, and 44.0%, and the three low-abundance species at 1.4%, 3.0%, and 3.0%, respectively (Supplementary Table S2).

There are at least three reasons for the remaining quantitative error. First, although we targeted 0.5x per reference skim, coverage still varied across species (Table S1), resulting in different powers of discrimination, as shown by the experiment with subsamples of *Knautia arvensis* (Supplementary Fig. S1). Fortunately, we found that even very low-depth skims of 0.05x are useful for species detection and are probably still useful for differentiating rare from abundant species (albeit with more error) (Supplementary Fig. S1). Genome sizes are also estimated with error, so it is also helpful that the subsampling experiment suggests that detection power asymptotes with higher sequencing depth (Supplementary Fig. S1), and as sequencing costs fall further, we expect that the most robust protocol will be to target 1x coverage.

Second, very closely related species can generate false positives. Our reference-skim database included six congener pairs, and we included two of the pairs (*Papaver* and *Ranunculus*) in the mock mixes. In the case of *Papaver*, there were no *P. rhoeas* false-positives greater than the 1% minimum-abundance filter in the mocks that contained *P. somniferum* (MM1 and MM6) (Supplementary Table S2). In contrast, *Ranunculus acris* was regularly incorrectly assigned to reads in mock mixes that contained the closely related congener *Ranunculus repens*. In fact, almost all the false-positive assignments (93.4%) were to *R. acris*. In retrospect, this result is expected because these two species are not easily differentiated by pollen morphology (Forup & Memmott, 2005), floral morphology, or even DNA barcodes (*rbcL* (99.1% similarity), *matK* (96.9%), *ITS2* (95.5%)). In other words, the RevMet results are correctly telling us that the two *Ranunculus* species are very closely related.

Third, MinION reads have relatively high error rates of roughly 5 to 10% depending on the flow cell and kit used (Leggett & Clark, 2017). Although this is dropping over time, this error rate unavoidably obscures differences between species (although not enough to confound the two *Papaver* species). We note that one of the advantages of the RevMet approach is that we use both sequence similarity and percent coverage as predictors of species presence (Fig. 1C). Using sequence similarity alone, we observed several instances of low numbers of mapped reads being given false-positive assignments (data not shown). The percent-coverage filter requires many reference-skim reads to independently identify a species before an assignment is made.

### Reference-skim cost

The RevMet pipeline is relatively low cost. In our study, we generated skims for 49 plant species, with genome sizes ranging from roughly 290 Mb (*Epilobium hirsutum*) to just under 15 Gb (*Sambucus nigra*), targeting 0.5x coverage. All skims were produced on a single lane of Illumina HiSeq 2500 (250 PE) at a mean coverage of 0.57x. The average cost per skim in this study was just under £90, which includes the DNA extraction, LITE library preparation, sequencing, and data QC. The per skim cost will be lower in studies with smaller eukaryotic genomes and should be lower with Illumina’s new sequencer, the NovaSeq 6000. Genome assembly projects are likely to produce many such datasets for free download in the future.

### Long-read MinION cost

We used ONT’s first iteration of the Rapid Barcoding Kit (RBK-001), which relies on transposase to randomly fragment DNA and simultaneously add barcoded adapters. Longer read lengths have an increased likelihood of accurate species assignment because they carry more sequence information. The two main ways to obtain longer reads with transposase-based preparations are to: (1) increase the ratio of DNA to transposase e.g. by increasing the input material or by heat killing a proportion of the transposase (which also lowers sequencing yields); and (2) use higher molecular weight input DNA. Since the release of RBK-001, ONT’s chemistry has evolved, and their Rapid-based kits have seen greater sequencing yields. However, the recommended input for the latest iteration of the Rapid Barcoding Kit (RBK-004) is now higher, 400 ng of DNA per sample. That said, we anticipate that input biomasses similar to those used in this study, 200 ng, will still be adequate. Also, even 400 ng is achievable, as 36 of 48 of our wild-bee pollen samples samples yielded >400 ng (Supplementary Table S3). ONT have also recently released the Flongle (∼$90), which is a disposable nanopore that allows prolonged reuse of the MinION. Our results suggest that ONT’s target yield of 1 Gb per Flongle will be more than enough for multiplexing twelve bee-collected pollen loads, reducing per-sample costs from the £61 in this study to just under £16.

### Application to pollen collected from wild bees

The RevMet pipeline detected consistent differences in the compositions of pollen loads collected by honeybees *Apis mellifera* and by the two bumblebees *Bombus terrestris/lucorum* and *B. lapidarius* (Fig. 3). Importantly, because our data are semi-quantitative, we are able to conclude that even the bumblebees showed fidelity to one plant species (Fig. 3C), a result that would be less reliably concluded from metabarcoding data.

The RevMet pipeline can readily be applied to a wide range of research questions. Most obviously, ReMet could be used to compare pollination networks across large-scale spatial and biogeographical gradients (Pornon *et al.,* 2016). RevMet could potentially also be used to quantify the degree to which co-attraction of pollinators leads not to benefits of increased pollinator numbers but to loss of pollination service via competition (Carvalheiro et al. 2014; Pornon *et al.,* 2016). Outside of pollination ecology, there is potential for semi-quantitative assessments of many other eukaryotic species mixtures, including herbivore diets (Bhattacharyya, Dawson, Hipperson, & Ishtiaq, 2018; Kress, García-Robledo, Uriarte, & Erickson, 2015); plant-fungus interactions (Schröter *et al.,* 2018); allergenic pollen species from air samples (Kraaijeveld *et al.,* 2015); and algal and diatom communities (Keller *et al.,* 2015). Furthermore, due to the portability and real-time nature of the MinION platform, the method could be optimised for analysis in the field alongside sample collection.

## Acknowledgements

This work was supported by the Norwich Research Park Science Links Seed Fund, the BBSRC Norwich Research Park Biosciences Doctoral Training Partnership (grant number BB/M011216/1) and BBSRC Core Strategic Programme Grant BB/CSP17270/1 to the Earlham Institute. LVD is funded by the Natural Environment Research Council (NE/N014472/1). CC was supported by the J. Arthur Ramsay Fund, administered by the Cambridge University Zoology Department. We are grateful for the support of the Earlham Institute Genomics Pipelines group and the NBI Computing infrastructure for Science (CiS) group. We thank Iain Barr for help in fieldwork and Pensthorpe Natural Park for access.

## Authors’ contributions

MDC, RML, DWY, LVD, and RGD conceived and designed the study. LVD, RGD, and CC collected the samples. NP, DH, and LP performed the experiments. NP, RML, and DWY analyzed the data. NP and DWY led the writing of the manuscript. All authors gave final approval for publication.

## Data accessibility

The Illumina and MinION datasets are available in the European Nucleotide Archive (http://www.ebi.ac.uk/ena) under study accession PRJEB30946. RevMet scripts are available from https://github.com/nedpeel/RevMet and a tutorial using an example dataset can be found at https://revmet.readthedocs.io/en/latest/.

## Figure legends and table titles

Figure S1. Numbers of mock-mix reads assigned to *Knautia arvensis*, and declines in the number of unassigned reads, at different reference-skim coverage levels. The subsampling was repeated three times.

Table S1. Estimated genome sizes, read counts, and coverage for the genome skim references.

Table S2. RevMet taxonomic assignments of mock-sample MinION reads.

Table S3. RevMet taxonomic assignments of bee-collected pollen MinION reads and pollen sample information.

