## Supplementary figures and images for "Semi-quantitative characterisation of mixed pollen samples using MinION sequencing and Reverse Metagenomics (RevMet)"

### Fig. S1

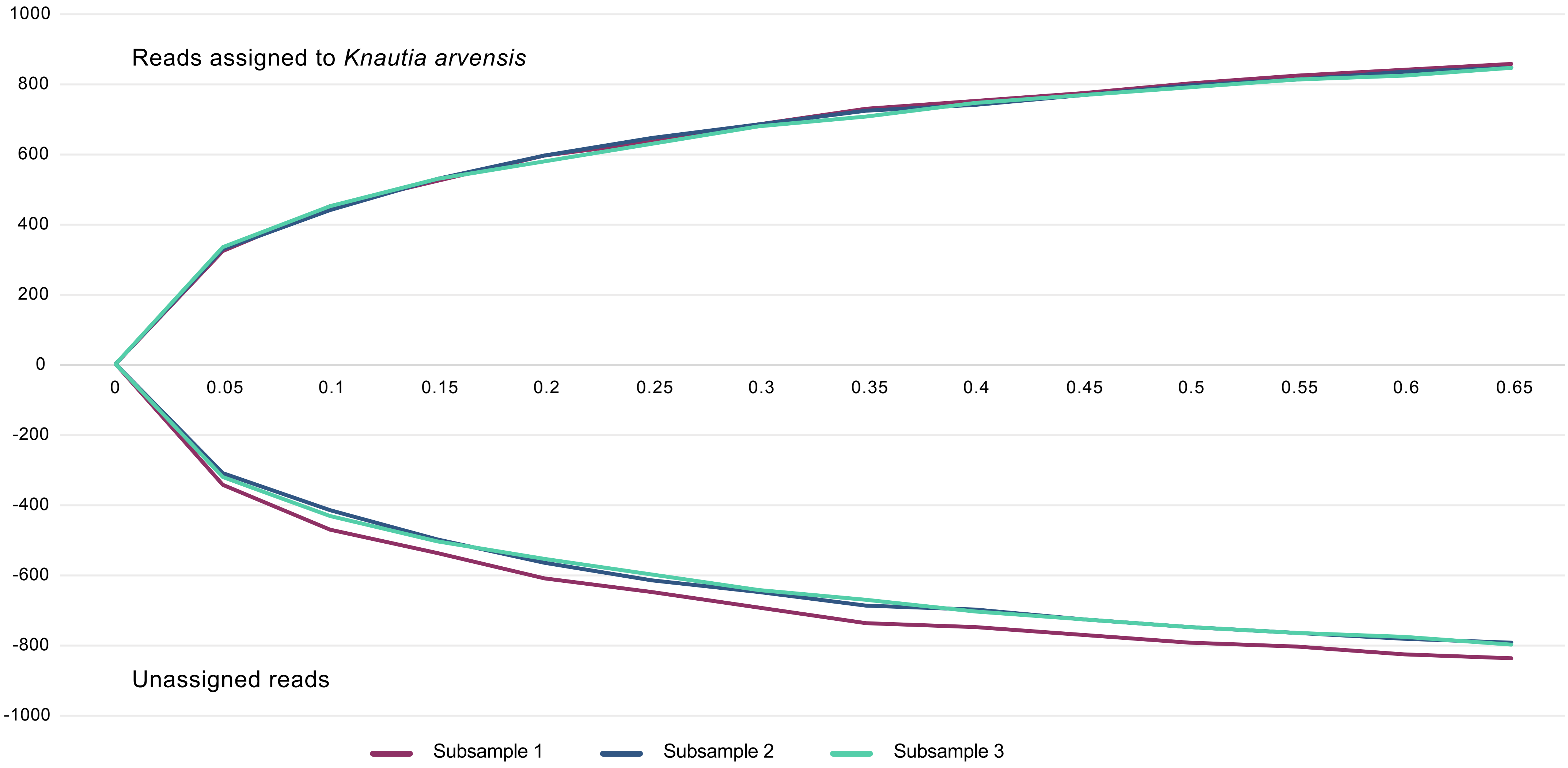
